# Discrimination among *Cryptococcus deneoformans, C. neoformans* and interspecies hybrids using MALDI-TOF Mass Spectrometry

**DOI:** 10.1101/2020.03.13.991661

**Authors:** Margarita Estreya Zvezdanova, Manuel J. Arroyo, Gema Méndez, Jesús Guinea, Luis Mancera, Patricia Muñoz, Belén Rodríguez-Sánchez, Pilar Escribano

**Author notes:** **Corresponding Author: Belén Rodríguez-Sánchez, PhD.** Servicio de Microbiología-Enfermedades Infecciosas. Hospital General Universitario Gregorio Marañón. Dr Esquerdo 46. 28007 Madrid, Spain. These authors have contributed equally to the study (Belén Rodríguez-Sánchez, Pilar Escribano).

## Abstract

**Background:** Differentiation of the species within the *Cryptococcus neoformans* complex (*C. deneoformans, C. neoformans* and *C. neoformans* interspecies hybrids –*C. deneoformans* x *C. neoformans*-) is important to define the epidemiology of the infection.

**Objectives:** In this study we attempted the discrimination of three *C. neoformans* species using MALDI-TOF MS coupled with an in-house library.

**Methods:** All *Cryptococcus* spp. isolates were identified by AFLP markers. An in-house database was constructed 26 well characterized *C. deneoformans, C. neoformans* and interspecies hybrids. Forty-four *Cryptococcus* spp. isolates were blindly identified using MALDI-TOF MS (Bruker Daltonics) and the expanded library. Their protein spectra were also submitted to hierarchical clustering and the resulting species were verified via Partial Least Squares Differential Analysis (PLS-DA) and Support-Vector Machine (SVM).

**Results:** MALDI-TOF MS coupled with the in-house library allowed 100% correct identification of *C. deneoformans* and *C. neoformans* but misidentified the interspecies hybrids. The same level of discrimination among *C. deneoformans* and *C. neoformans* was achieved applying SVM. The application of the PLS-DA and SVM algorithms in a two-step analysis allowed 96.95% and 96.55% correct discrimination of *C. neoformans* from the interspecies hybrids, respectively. Besides, PCA analysis prior to SVM provided 98.45% correct discrimination of the 3 species analysed in a one-step analysis.

**Conclusions:** Our results indicate that MALDI-TOF MS could be a rapid and reliable tool for the correct discrimination of *C. deneoformans* and *C. neoformans*. The correct identification of the interspecies hybrids could only be achieved by hierarchical clustering with other protein spectra from the same species.

## INTRODUCTION

The genus *Cryptococcus* has classically comprised two sibling species with great importance from the clinical point of view: *Cryptococcus neoformans* and *C. gattii*, the causative agents of cryptococcosis. Whilst *C. neoformans* complex has been associated with meningitis in immunosuppressed patients, *C. gatti* has been shown to cause disease in both immune competent and immunocompromised population^1,2^. Species differentiation is important in order to establish the epidemiology, virulence and susceptibility pattern to the commonly used antifungal drugs^3-6^. So far, species assignment is achieved by morphology analysis of the colonies grown on specific culture media and serological tests^7^. The availability of DNA-based methodologies as restriction fragment length polymorphism (RFLP) analysis^8^, amplified fragment length polymorphism (AFLP) analysis^9^, multilocus microsatellite typing -MLMT-^10^, and multilocus sequence typing –MLST-^11^ has allowed the identification of *Cryptococcus* species and molecular types in the last years^8-13^. Genotyping methods have identified the following major molecular types: AFLP1/VNI, AFLP1A, AFLP1B/VNII for *C. neoformans*; AFLP2/VNIV for *C. deneoformans*, AFLP3/VNIII for the interspecies hybrid *C. neoformans neoformans* x *C. deneoformans*; and AFLP4/VGI, AFLP5/VGIII, AFLP6/VGII, AFLP7/VGIV and AFLP10/VGIV, VGII for *C. gattii* compex^14,15^.

Molecular techniques have shown to be accurate and robust although the whole procedure is cumbersome, time consuming, and delays the final identification. Although genomic analysis is currently the gold standard for *Cryptococcus* identification, its high requirements in hands-on time and expertise has led to the evaluation of alternative tools.

Matrix-assisted laser desorption ionization–time of flight mass spectrometry (MALDI-TOF MS) has emerged as a promising technology for the rapid and reliable identification of yeasts^16-18^. Isolates belonging to the *Candida* genus have been shown to be easily identified at the species level either from single colonies or directly from clinical samples using MALDI-TOF MS^19^. However, non-Candida yeasts still represent a challenge for this technology, especially when trying to identify genera poorly represented or even lacking in the commercial databases^20^. In this case, expanded in-house databases containing protein spectra from the underrepresented species and genera have shown to overcome this drawback^16^. Although this approach has worked before for the discrimination between *C. neoformans* and *C. gatti* complexes^21,22^, the available information about MALDI-TOF discrimination within the *C. neoformans* complex is still limited^23^.

In this study, MALDI-TOF has been applied for the discrimination between *C. deneoformans, C. neoformans* and the interspecies hybrids. For this purpose, two approaches have been applied: a database was built using well-characterized isolates and automated peak analysis was performed.

## MATERIALS AND METHODS

### Isolates and molecular identification

We retrospectively selected 70 *Cryptococcus* spp. isolates from clinical samples (n=70) belonging to 67 patients admitted to Hospital Gregorio Marañón (Madrid, Spain) from 1994 to 2007. Isolates sourced from cerebrum spinal fluids (51%), blood (33%), respiratory samples (10%), and others (6%). They were morphologically identified on Columbia agar + 5% sheep blood plates (Biomérieux Marcy L’étoile, France) at 35 ° C, and by means of the ID 32C system (bioMérieux, Marcy l’Etoile, France). All isolates were stored at −80°C in water until further analysis. All isolates were previously identified by AFLP analysis^24^ and were stored at −80°C in water until further analysis. Molecular identifications were considered as the reference in our study.

### Database construction

Twenty-six *Cryptococcus* isolates - *C. neoformans* (n=12), interspecies hybrids (n=10) and *C. deneoformans* (n=4)- were processed according to the manufacturer’s instructions and added to the in-house database (HGM library) as individual Main Spectra (MSPs).

The procedure for adding new entries to an in-house library has already been described^25^. Briefly, the instrument was calibrated before spectra acquisition using freshly prepared BTS; *Cryptococcus* isolates were processed as explained below and then spotted onto eight positions in the MALDI target plate and each position was read three times. Twenty-four protein spectra were thus achieved, 20 of which had to be identical in order to be accepted by the software (Biotyper, Bruker Daltonics, Bremen, Germany) as a MSP and added to the extended library.

### MALDI-TOF identification

Forty-four *Cryptococcus* spp. isolates were blindly analysed using an LT Microflex benchtop MALDI-TOF mass spectrometer (Bruker Daltonics) for spectra acquisition, using default settings. For the identification of the protein spectra, the updated BDAL database containing 8223 MSPs (Bruker Daltonics) was applied. This database contains 12 reference MSPs from *C. neoformans* and 7 from *C. deneoformans*. Besides, the expanded in-house HGM library developed in this study was used in combination with the commercial database.

The sample processing method applied consisted of a mechanical disruption step followed by a standard protein extraction. Briefly, a few colonies were picked, re-suspended in 300 μl water HPLC-grade and 900μl ethanol, and submitted to 5 min vortexing. After a brief spin, the supernatant was discarded and the pellet allowed drying completely at RT. Protein extraction with formic acid and acetonitrile was performed and 1μl of the supernatant was spotted onto the MALDI target plate in duplicates. Once the spots were dry, they were covered with 1μl HCCA matrix (Bruker Daltonics), prepared following the manufacturer’s instructions (Figure 1).

**Figure 1.**
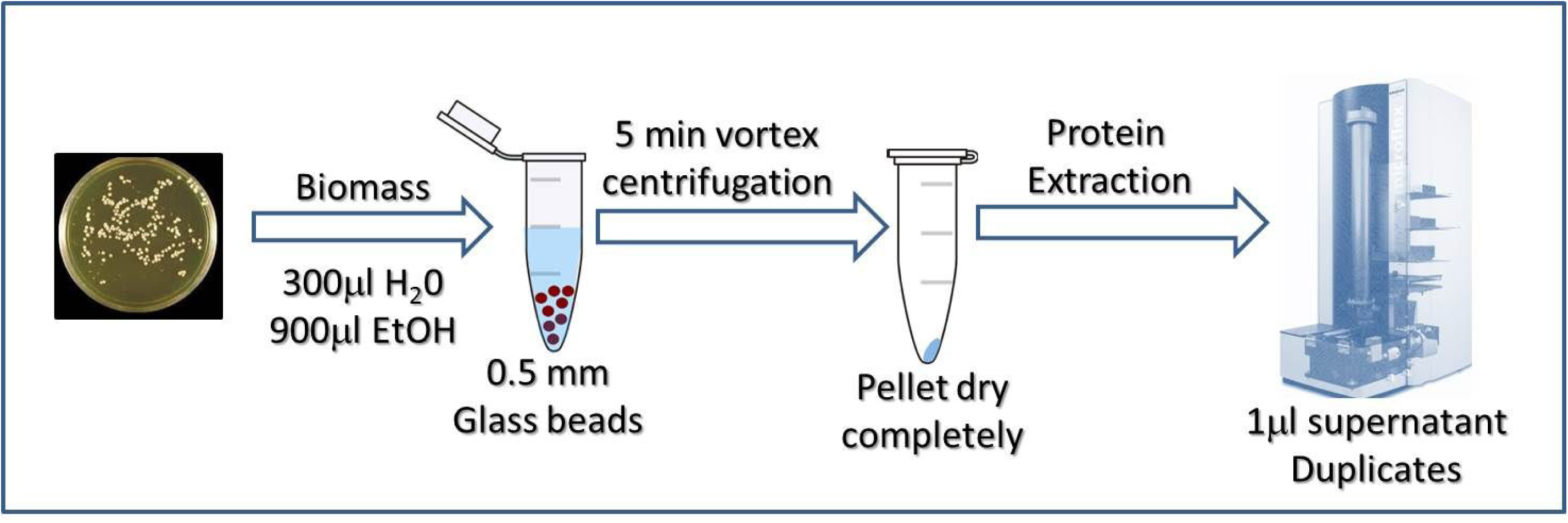
Workflow of the sample preparation method used in this study to obtain proteins from *Cryptococcus* spp. isolates for their identification by MALDI-TOF MS.

The identifications provided by MALDI-TOF MS were compared at the species level with those provided by AFLP analysis regardless of their score value (Table 1). Besides, score values ≥2.0 were considered as “high-confidence” scores and those ≥1.7 as “low-confidence” ones. Score values below 1.6 were only considered when consistent over the four top identifications, otherwise they were considered as “not reliable”.

**Table 1.**
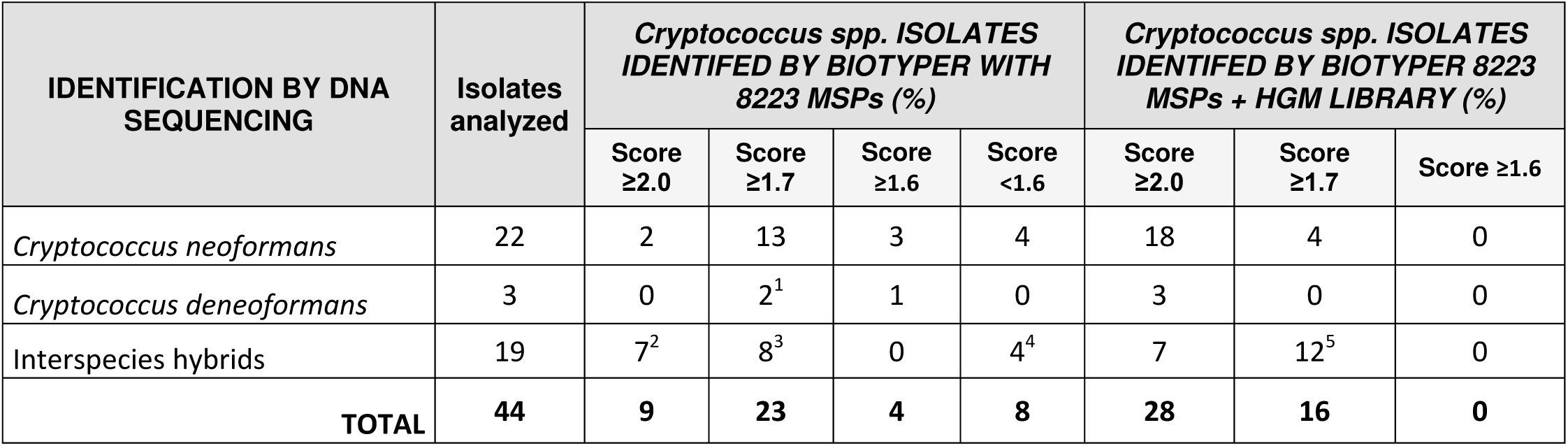
Identification of *Cryptococcus neoformans, C. deneoformans* and interspecies hybrids using MALDI-TOF MS and the Biotyper library alone or in combination with the in-house HGM database. ^1^Identified as *C. neoformans* complex (n=2); ^2^Identified as *C. neoformans* complex (n=7); ^3^Identified as *C. neoformans* complex (n=1), *C. deneoformans* (n=4) and *C. neoformans* (n=3); ^4^Identified as *C. neoformans* complex (n=1) and *C. deneoformans* (n=3); 5Identified as *C. neoformans* (n=7)

### Peak Analysis

For the classification of the three species of *Cryptococcus* their protein spectra were processed using Clover MS Data Analysis software (Clover Biosoft, Granada, Spain) with the parameters shown in Table S1 in order to achieve a peak matrix with a representative mass list in the range 2400m/z to 12000m/z. Furthermore, spectra alignment was performed. First, the replicates from the same isolate were aligned in order to get an average spectrum. Finally, all average spectra were aligned together.

The rate of presence for the biomarker peaks was calculated for each species and then compared among species. Receiver Operating Characteristic (ROC) curve with Area under the Curve –AUC- higher than 0.99 were used as quality indicators to measure the sensibility and specificity of a selected biomarker.

Once the putative biomarkers were selected and analysed, a peak matrix was built containing all the aligned spectra from all *Cryptococcus* isolates, processed as described in Table S2. This peak matrix was constructed with ten species-specific biomarkers and it was used as input for a dendrogram obtained measuring Euclidean distance from Principal Component Analysis (PCA) scores.

Over the peak matrix, two approaches were applied in order to discriminate the three *Cryptococcus* species. The first one was a two-step method in which the discrimination of *C. deneoformans* from the other two species was performed as a first step and it was replicated by means of two machine learning algorithms on the same peak matrix: supervised PLS-DA and SVM. Results were validated using k-fold cross validation method. In the second step, a new peak matrix was performed in order to achieve a better discrimination of *C. neoformans* from the interspecies hybrids. A second dendrogram was performed using the above mentioned parameters. Again, PLS-DA and SVM was performed to this second peak matrix to replicate the classification and the k-fold cross validation method was applied. The two-step method was further improved by the exclusion from the peak matrix of peaks that did not provide enough discrimination.

In order to simplify the workflow, a one-step method was assayed so that the capacity of the algorithms to discriminate the three *Cryptococcus* species at the same time was tested. In this case, only one peak matrix with spectra from the three species was built and 5 species-specific biomarkers were included. The alignment and processing parameters were the same as in the two-steps approach. The one-step method was evaluated using the peak matrix generated as input data for PLS-DA analysis and SVM analysis. Besides, the validation in both cases was performed using k-fold confusion matrix.

### Ethic Statement

The hospital Ethics Committee approved this study and gave consent for its performance (Code: MICRO.HGUGM.2017-003). Since only microbiological samples were analysed, not human products, all the conditions to waive the informed consent have been met.

## RESULTS

Genotyping of the isolates detected three different genotypes. The most common genotype was AFLP1/1B (*C. neoformans*, n=34; 49%), followed by AFLP3 (interspecies hybrids, n=29; 41%) and AFLP2 (*C. deneoformans*, n=7; 10%).

The application of MALDI-TOF MS and the commercial database allowed the correct identification of 18/22 *C. neoformans* isolates (81.8%) and 1/3 *C. deneoformans* isolates (33.3%); the remaining *C. neoformans* isolates –n=4- could not be reliably identified and for 2 *C. deneoformans* isolates MALDI-TOF did not provide the species identification (Table 1). The identification of the interspecies hybrids (n=19) was not achieved using the commercial database due to the lack of representation of this microorganism. These isolates were identified as *C. neoformans* complex in 9 cases (score≥2.0, n=7; score>1.7, n=1; score<1.6, n=1), as *C. deneoformans* in 7 cases (score>1.7, n=4; score<1.6, n=3) and as *C. neoformans* in 3 cases (score>1.7) –Table 1-.

Only two isolates (8.0%) were correctly identified at the species level with high-confidence score values (≥2.0) whilst 52.3% of the samples were identified with low-confidence scores (>1.7) - Table 1-. Another 4 isolates were reliably identified to the species level, although with scores values ranging between 1.7 and 1.6 and, finally, 8 isolates obtained scores below 1.6. The latter can be considered as unreliable identifications.

Using the in-house library all *C. neoformans* and *C. deneoformans* isolates were correctly identified by MALDI-TOF MS at the species level (Table 1). Moreover, 21/25 isolates (84.0%) were identified with score values ≥2.0 which indicates a high-confidence level. The reliability of the identification was further demonstrated by the fact that the top 4-5 identifications were identical in all cases. In all but two cases these top reference isolates belonged to the HGM in-house library.

However, the implementation of the expanded HGM library only allowed the correct identification of 12/19 interspecies hybrids, 7 of them with score values above 2.0. The high closeness of the interspecies hybrids with the other two *Cryptococcus* species made it difficult for MALDI-TOF MS to discriminate among them and misidentified 7 interspecies hybrids as *C. neoformans* (Table 1).

To improve the identification of the interspecies hybrids and their discrimination from *C. deneoformans* and *C. neoformans*, peak analysis was performed. The search for species-specific biomarker peaks yielded a list of 10 peaks that allowed the differentiation of the *Cryptococcus* species analysed, with 5 of them showing higher discriminative power (Table 2). The two-step method allowed correct differentiation of the interspecies hybrids which clustered distinctly in the dendrograms built using two different hierarchical clustering variations (Figure 2 and Figure S2). These dendrograms showed three different clusters where *Cryptococcus deneoformans* isolates were clearly separated from *Cryptococcus neoformans* and the interspecies hybrids. Accurate differentiation among the 3 *Cryptococcus* species was achieved using the peak matrix built upon the 5 most discriminative peaks, with only one spectrum from an interspecies hybrid misallocated in the *C. neoformans* cluster (Figure 2B). *C. neoformans* and the interspecies hybrids showed close relatedness between them based on their protein spectra.

**Table 2.** List of the 10 representative mass peaks of *Cryptococcus* spp. Identified as potential biomarkers. These peaks were used for the construction of dendrograms and PLS-DA and SVM models. The 5 peaks marked with asterisks (*) were selected for the simplified models. CV= Coefficient of Variation.

| Mass (m/z) | Number of Spectra | *C. neoformans* | *C. neoformans* (CV) | *C. neoformans* (mean) | Interspecies hybrids | Interspecies hybrids (CV) | Interspecies hybrids (mean) | *C. deneoformans* | *C. deneoformans* (CV) | *C. deneoformans* (mean) |
| --- | --- | --- | --- | --- | --- | --- | --- | --- | --- | --- |
| 2488.07 | 54 | 30/34 | 88.749 % | 4.401.767 | 24/24 | 69.617 % | 2.811.789 | 0/7 | - | - |
| 2842.14 | 53 | 29/34 | 78.206 % | 2.202.128 | 24/24 | 79.602 % | 2.235.196 | 0/7 | - | - |
| *3084.11 | 55 | 31/34 | 98.458 % | 7800.06 | 24/24 | 89.283 % | 5.969.393 | 0/7 | - | - |
| *5453.91 | 27 | 1/34 | 0.0 % | 72.906 | 23/24 | 65.081 % | 731.902 | 3/7 | 12.654 % | 748.872 |
| *5552.90 | 27 | 1/34 | 0.0 % | 558.307 | 23/24 | 73.624 % | 1.418.905 | 3/7 | 47.3 % | 2.763.278 |
| 6576.08 | 23 | 0/34 | - | - | 16/24 | 63.172 % | 457.978 | 7/7 | 56.698 % | 685.58 |
| *6688.67 | 57 | 34/34 | 95.69 % | 4.420.907 | 23/24 | 88.388 % | 3.556.217 | 0/7 | - | - |
| *7103.01 | 31 | 1/34 | 0.0 % | 24.32 | 23/24 | 122.759 % | 1.484.767 | 7/7 | 52.14 % | 4.155.275 |
| 7830.42 | 18 | 0/34 | - | - | 11/24 | 46.494 % | 719.13 | 7/7 | 39.831 % | 449.704 |
| 8636.24 | 43 | 19/34 | 101.061 % | 2.887.856 | 24/24 | 87.315 % | 1.832.722 | 0/7 | - | - |

**Figure 2.**
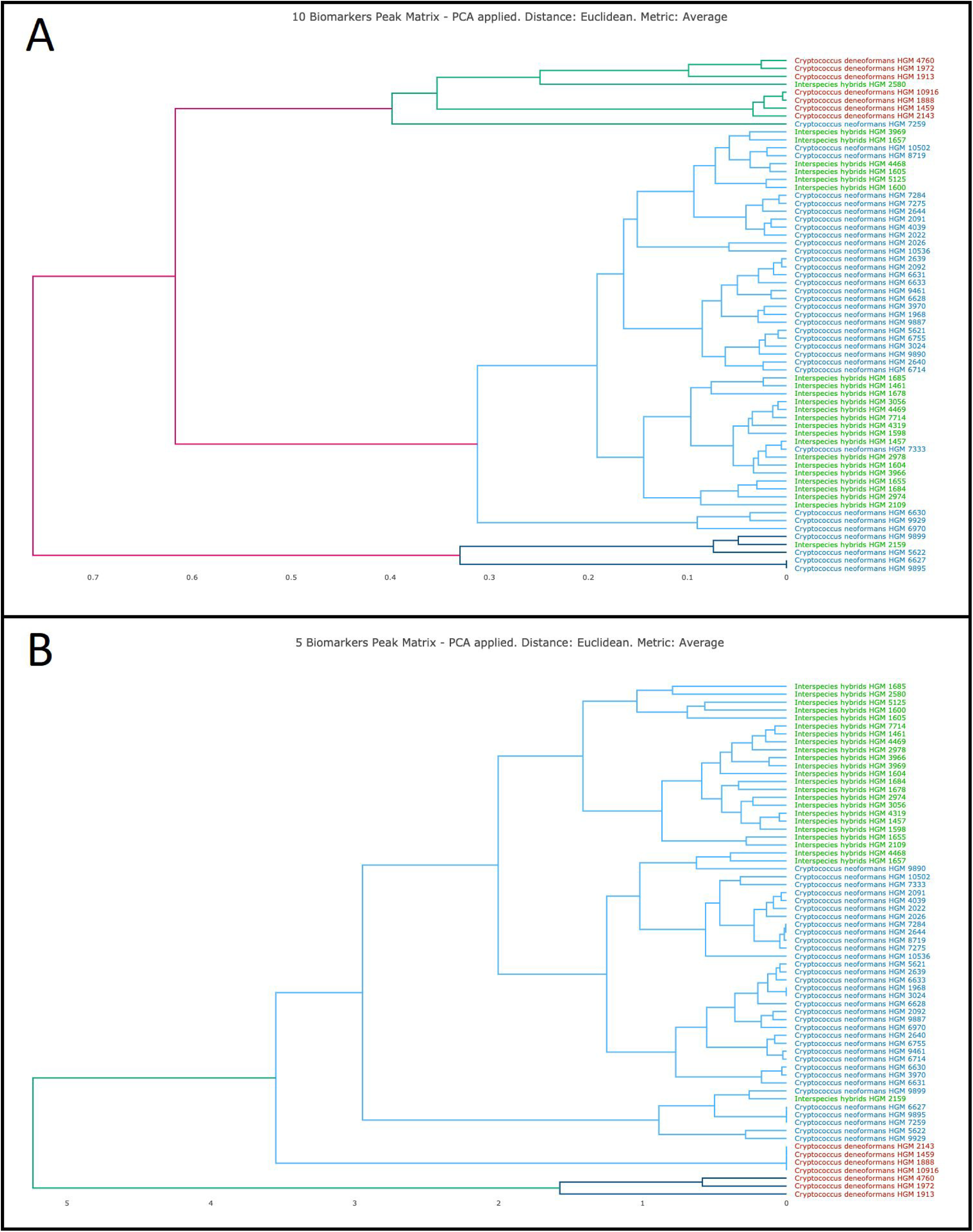
Clustering of 65 *Cryptococcus* isolates included in this study. Five isolates could not be recovered from culture for further analysis.

The validation of the method yielded a k-fold (k=10) score of 96.92% for PLS-DA performed over the peak matrix with 10 biomarkers and 98.46% for the analysis with 5 biomarkers. However, SVM algorithm achieved 100% discrimination in both cases when PCA was applied (Table S3).

A second dendrogram was performed using hierarchical clustering analysis. It showed two well-defined clusters for *Cryptococcus neoformans* and the interspecies hybrids (Figure S2). In this step only the 3 biomarkers to differentiate *C. deneoformans* from interspecies hybrids were used (5453.91, 5552.90 and 7103.00 m/z). Furthermore, this second dendrogram was validated by PLS-DA and SVM algorithms. K-fold (k=10) was applied achieving 95.55% efficacy in both analyses.

In the single-step method, the peak matrix built with 5 biomarkers was used as an input for PLS-DA and SVM analysis in order to achieve the discrimination of the 3 *Cryptococcus* species simultaneously. PLS-DA analysis could not classify correctly the three varieties at the same time due to the low k-fold (k=10) values obtained. However, PCA performance prior to SVM allowed 98.46% correct classification of the three *Cryptococcus* species (Figure 3). The efficacy of the method was tested by k-fold (k=10) cross validation analysis was above 95.0%. (Figure 3, Table S3)

**Figure 3.**
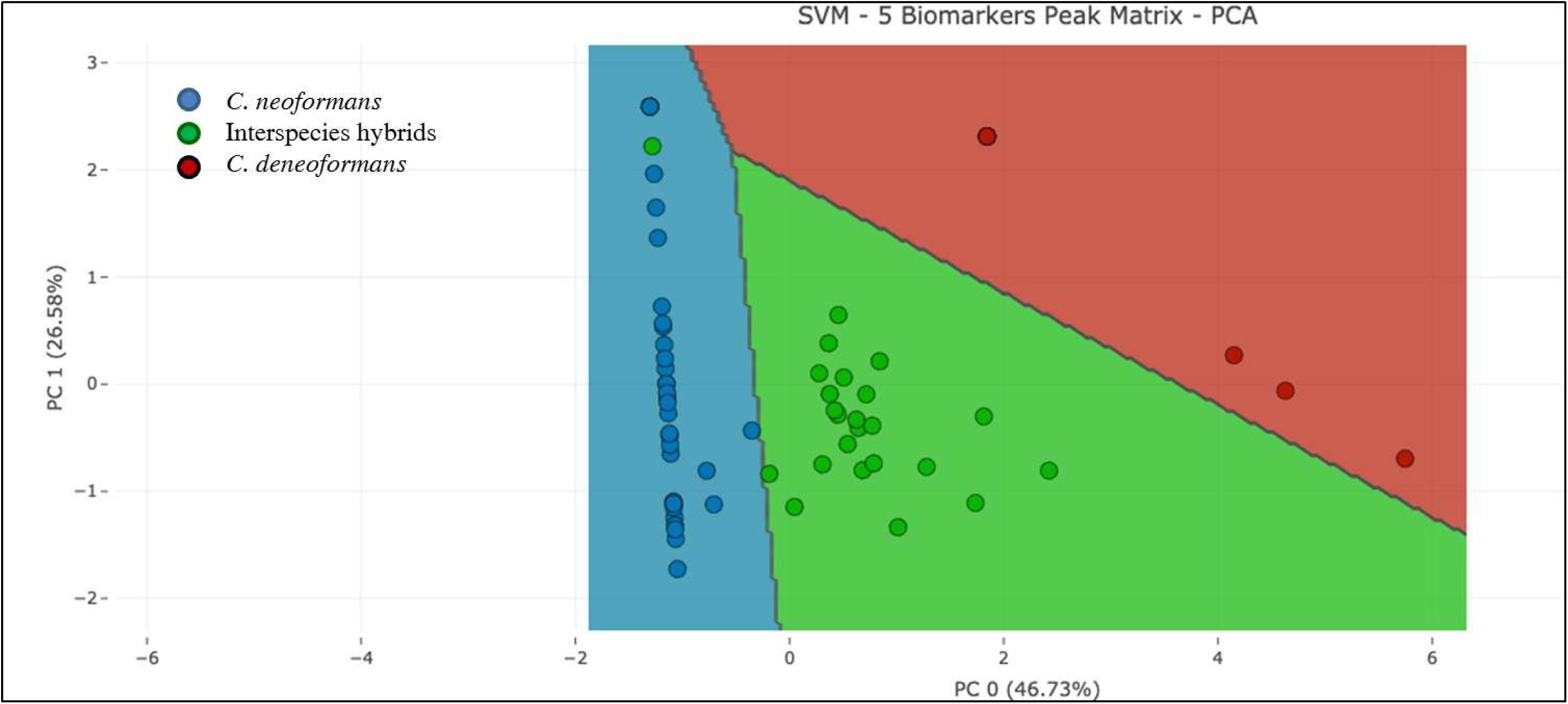
Classification of the three *Cryptococcus* species by SVM in the one-step approach, using 5 biomarker peaks.

As a result of this analysis, a visual method for the differentiation of the analyzed *Cryptococcus* species can be applied based on the presence of the 6688.67 m/z peak in the *C. neoformans* isolates and their absence in *C. deneoformans* isolates, where the peaks 6576.08 m/z and 7103.01 m/z could be detected. On the other hand, both sets of peaks are present in the interspecies hybrids although some of them (2842.14, 3084.11 and 8636.24 m/z) were detected in 100% of the spectra from this species (Table 3). The visual detection of these biomarker peaks could provide a rapid and accurate identification of the *Cryptococcus* species prior to a more in-depth peak analysis using *ad-hoc* software.

**Table 3.** Differentiation of the analyzed *Cryptococcus* species based on the absence/presence of biomarker peaks. Figures indicate the percentage (%) of isolates showing the indicated peak.

| m/z | 2842.14 | 3084.11 | 6576.08 | 6688.67 | 7103.01 | 8636.24 |
| --- | --- | --- | --- | --- | --- | --- |
| *C. deneoformans* | 0 | 0 | 100 | 0 | 100 | 0 |
| *C. neoformans* | 85.3 | 91.2 | 0 | 100 | 0 | 55.9 |
| *Interspecies hybrids* | 100 | 100 | 66.7 | 95.8 | 4.3 | 100 |

## DISCUSSION

Accurate identification of *Cryptococcus* species within the *Cryptococcus neoformans* complex provides valuable information about their epidemiology, sensitivity to commonly used antifungal drugs or virulence. Our results show that discrimination among the three *Cryptococcus* species analyzed–*C. deneoformans, C. neoformans* and interspecies hybrids-can be performed successfully using MALDI-TOF MS.

The implementation of the in-house database built in our laboratory allowed 100% correct species-level identification of the 25 *Cryptococcus deneoformans* and *C. neoformans* isolates used to challenge it. Apart from the reliable identification of the analyzed *Cryptococcus* species, the in-house library also provided high confidence identifications in 63.6% of the cases (Table 1). Furthermore, these results showed consistency along the 10 top identifications provided by the mass spectrometry instrument, even for the hybrids. This fact is of great importance in the routine of the microbiology laboratory in order to transfer reliable information to the clinicians.

The results obtained are in agreement with those obtained by other authors^21-23^. However, the in-house library did not provide enough discrimination between the above-mentioned species and the interspecies hybrids. This goal was only fulfilled completely when peak analysis was performed and the three *Cryptococcus* species analyzed in this study distinctively clustered together (Figure 2). Other authors have provided species-level discrimination in 98.1-100% of the cases^21, 23, 26^. Although some of these studies were performed on higher number of isolates, our results also reflect the improvements made on the commercial database during the last years.

The available commercial database has demonstrated to provide high species-level resolution for *C. deneoformans* and *C. neoformans*–76.0%- although score values <1.7 were obtained in 21.0% of the cases and species-level identification was not provided for 2 *C. deneoformans* isolates. These data supported the need of building expanded databases. However, even improvements in the reference databases proved not to be enough to differentiate the interspecies hybrids. This may be due to the algorithms used by the mass spectrometry instrument for species assignment and to the fact that the hybrids show peaks present of both parental species. Therefore, peak analysis using *ad-hoc* software was performed. A list of 10 biomarker peaks was achieved as the input for species classification (Table 2). The implementation of PLS-DA analysis in a two-step approach allowed the discrimination of *C. deneoformans* isolates in the first place and, subsequently, the correct classification of *C. neoformans* isolates and the interspecies hybrids in 96.92% of the cases. Furthermore, the accuracy of this method increased when the number of biomarker peaks used was reduced to the five most discriminative ones (98.46%).

In order to simply the analysis, a one-step method was proposed in order to classify the three species simultaneously. In this case, PLS-DA provided correct classification in less than 75.0% of the cases but the application of SVM after PCA analysis allowed 96.92% correct discrimination of the analyzed isolates. This analysis provided a set of species-specific peaks for the *Cryptococcus* species within the *C. neoformans* complex that may be detected by visual inspection, representing a rapid and inexpensive approach for their discrimination.

In summary, our results demonstrate the usefulness of MALDI-TOF MS when applied in the microbiology laboratory for rapid and reliable identification of non-*Candida* yeasts. Although the updated commercial library provided correct species-level identification for a high number of *C. deneoformans* and *C. neoformans* isolates (43.2%), the identification of these species was missing or not reliable in 20.5% 18.2% of the cases, respectively. Moreover, the detection of the interspecies hybrids is not possible with the Biotyper database. However, the expanded in-house library allowed correct species-level identification for all *C. deneoformans* and *C. neoformans*, either by conventional identification with MALDI-TOF MS or by peak analysis (Figure 3). The interspecies hybrids required hierarchical clustering for their correct identification since their close relatedness with the other species made it difficult for MALDI-TOF to differentiate them from the other two species in a routine manner. This approach and the detection of species-specific peaks are recommended for the reliable discrimination of the three analyzed species.

## Acknowledgements

The authors want to thank Álvaro Gómez-González for his assistance with Bruker Daltonics databases. This study has been supported by the Miguel Servet Program (ISCIII-MICINN CP14/00220) and by the projects PI16/01012 (PE) and PI18/00997 (BRS) from the Health Research Fund (FIS) of the Carlos III Health Institute (ISCIII), Madrid, Spain, partially financed by the by the European Regional Development Fund (FEDER) ‘A way of making Europe.’ BRS (CPII19/00002), PE (CPI15/00115) and JG (CPII15/00006) are recipients of a Miguel Servet contract supported by the FIS program. The funders had no role in the study design, data collection and analysis, decision to publish, or preparation of the manuscript.

